# Antisense oligonucleotide targeting pathogenic sense repeat RNA in *C9ORF72* suppresses production of antisense-dependent dipeptide repeat proteins implicated in ALS/FTD

**DOI:** 10.1101/2024.10.04.616663

**Authors:** Yuanzheng Gu, Mark W. Kankel, Jonathan Watts, Paymaan Jafar-nejad, Sandra Almeida

## Abstract

A six nucleotide repeat expansion in intron-1 of the *C9ORF72* gene is the most common genetic mutation affecting individuals with Amyotrophic Lateral Sclerosis and Frontotemporal Dementia. Bi-directional transcription of the repeat expansion generates sense and antisense repeat RNAs that can then be translated in all reading frames to produce six distinct dipeptide repeat (DPR) proteins with unique termini. The precise site of translation initiation of these proteins within the *C9ORF72* repeat expansion remains elusive. We used CRISPR-Cas9 genome editing and steric-blocking antisense oligonucleotides (ASOs) to investigate the contribution of different AUG codons in the antisense repeat RNA to the production of DPR proteins, poly(GP) and poly(PR) in *C9ORF72* expansion carrier motor neurons and lymphoblast cells. We then utilized ASOs targeting *C9ORF72* sense repeat RNA to examine whether sense or antisense RNA is the major source of the poly(GP) protein - a question for which conflicting evidence exists. We found that these ASOs reduced the intended sense RNA target, but also the antisense RNA, thus preventing the production of poly(PR). Our data highlights the importance of the sequences preceding the antisense CCCCGG repeat expansion for the synthesis of antisense DPR proteins and supports the use of sense *C9ORF72* ASOs to prevent the accumulation of both sense- and antisense-dependent DPR proteins in *C9ORF72* ALS/FTD.

## Introduction

Amyotrophic lateral sclerosis (ALS) and Frontotemporal dementia (FTD) are two neurodegenerative disorders frequently caused by a GGGGCC (G_4_C_2_) repeat expansion in the *C9ORF72* gene^8,20^, a condition termed *C9ORF72* ALS/FTD. The mutation drives pathogenesis through a combination of loss of C9ORF72 protein normal function and gain of toxic effects due to the repeat expansion, which is bidirectionally transcribed to generate sense and antisense RNA transcripts. These repeat-containing sense and antisense RNAs can each be translated in all three reading frames to produce dipeptide repeat (DPR) proteins: poly(GA), poly(GR) and poly(GP) from sense RNA, and poly(GP), poly(PA) and poly(PR) from antisense RNA. Each expanded repeat RNA strand is commonly observed in RNA foci in human autopsy tissues along with the DPR proteins, both of which are considered pathological hallmarks of *C9ORF72* ALS/FTD disease^7^.

Overexpression of DPR proteins in cellular and animal models indicates that these proteins are highly toxic and impact many cellular functions^2^. Thus, significant attention has focused on identifying specific processes involved in the generation of these proteins to enable the development of prophylactic strategies to block their production. Despite intensive effort, it is unclear how the DPR proteins come to be synthesized from the *C9ORF72* RNAs harboring repeat expansions in *C9ORF72* hexanucleotide repeat expansion (HRE) carrier cells.

Multiple translation mechanisms likely play a role in the production of DPR proteins, influenced by factors such as the precise sequence context, cell type and stress conditions. The presence of initiation codons in the RNA sequence preceding the repeat expansion appears to be a key element, similar to the mechanism described for fragile X mental retardation 1 (FMR1) RNA^13,22^. In the *C9ORF72* RNA, a CUG near-cognate initiation codon 24 nucleotides upstream of the G_4_C_2_ repeats in the poly(GA) reading frame is required for poly(GA) synthesis in reporter constructs^4,10,23,26^. The necessity of this CUG was confirmed in the endogenous context in human motor neurons (MNs) carrying more than 1,000 G_4_C_2_ repeats where deletion of an intronic region including the CUG initiation site eliminated poly(GA) protein production^1^. The effect was selective, as the levels of the other two sense RNA reading frame DPR proteins, poly(GP) and poly(GR), were unaffected by this deletion. While repeat expansion could potentially drive positioning of the ribosomes in the intron, ribosome profiling studies reveal that elimination of the repeat expansion does not remove the ribosome footprints on the CUG codon used to initiate translation of the poly(GA)^29^. Collectively, these studies demonstrate that poly(GA) protein production relies on the RNA sequence upstream of the G_4_C_2_ repeat expansion. Poly(GR) protein production appears to follow a near cognate initiation mechanism like poly(GA), as deletion of 111 nucleotides upstream of the G_4_C_2_ repeats largely decreased translation in the poly(GR) frame, whereas removing only the repeats themselves did not affect translation levels as assessed with a luciferase reporter^15^.

The importance of the precise RNA sequence preceding the repeat expansion is greater for poly(GP) and poly(PR), as evidenced by the presence of canonical AUG initiation codons in the antisense RNA reading frames for both DPR proteins^30^. A study using a reporter system found that at least one of the three AUG codons in the poly(GP) frame is needed for its production^4^, an observation that was recently found to be reproducible^24^. Though poly(GP) is potentially made from the sense RNA, an analysis of human autopsy tissues from *C9ORF72* expansion carriers showed that the poly(GP) protein is predominantly generated using antisense RNA^31^. Nonetheless, antisense oligonucleotides (ASOs) that target the sense *C9ORF72* RNA greatly reduce poly(GP) protein in cells and cerebrospinal fluid from expansion carriers^9,27^, indicating that poly(GP) is produced from the sense RNA.

Here, we used CRISPR-Cas9 genome editing and steric-blocking ASOs to investigate the contribution of the region containing the AUG codons in the antisense repeat RNA to the production of poly(GP) and poly(PR) proteins in *C9ORF72* HRE carrier MNs and lymphoblast cells. We then characterized RNase-H-sensitive ASOs targeting the *C9ORF72* sense repeat RNA in an attempt to reconcile the conflicting evidence for whether sense or antisense RNA is the major source of the poly(GP) protein.

## Results

### Deletion of the AUG codons region in the antisense repeat RNA greatly reduces poly(GP) protein levels in *C9ORF72* iPSCs and motor neurons

Poly(GP) is the only DPR that can potentially be generated from both sense and antisense strands of the repeat RNA. In the sense direction, a stop codon located immediately upstream of the G_4_C_2_ repeats in the poly(GP) frame may prevent its translation. However, in the antisense (AS) direction, three AUG start codons are in frame with poly(GP). Unlike the sense strand, the AS strand is devoid of a stop codon to prevent translation into the C_4_G_2_ repeats. To determine the role of these AUG start codons, we used CRISPR-Cas9 technology to generate a homozygous deletion (AS-AUGs deletion) 3’ to the G_4_C_2_ repeats in iPSC lines from two *C9ORF72* HRE carriers (Fig. 1a). The deletion eliminated a 181-nucleotide sequence covering the three AUG codons in the poly(GP) frame and one AUG codon in the poly(PR) frame of the AS repeat RNA (Fig. 1b). No alternative start site in the poly(GP) or poly(PR) frames in the remaining sequence 5’ to the C_4_G_2_ repeats was created by the deletion. We tested other combinations of guide RNAs with the CRISPR-Cas9 in an attempt to generate smaller deletions but were unsuccessful.

**Fig 1.**
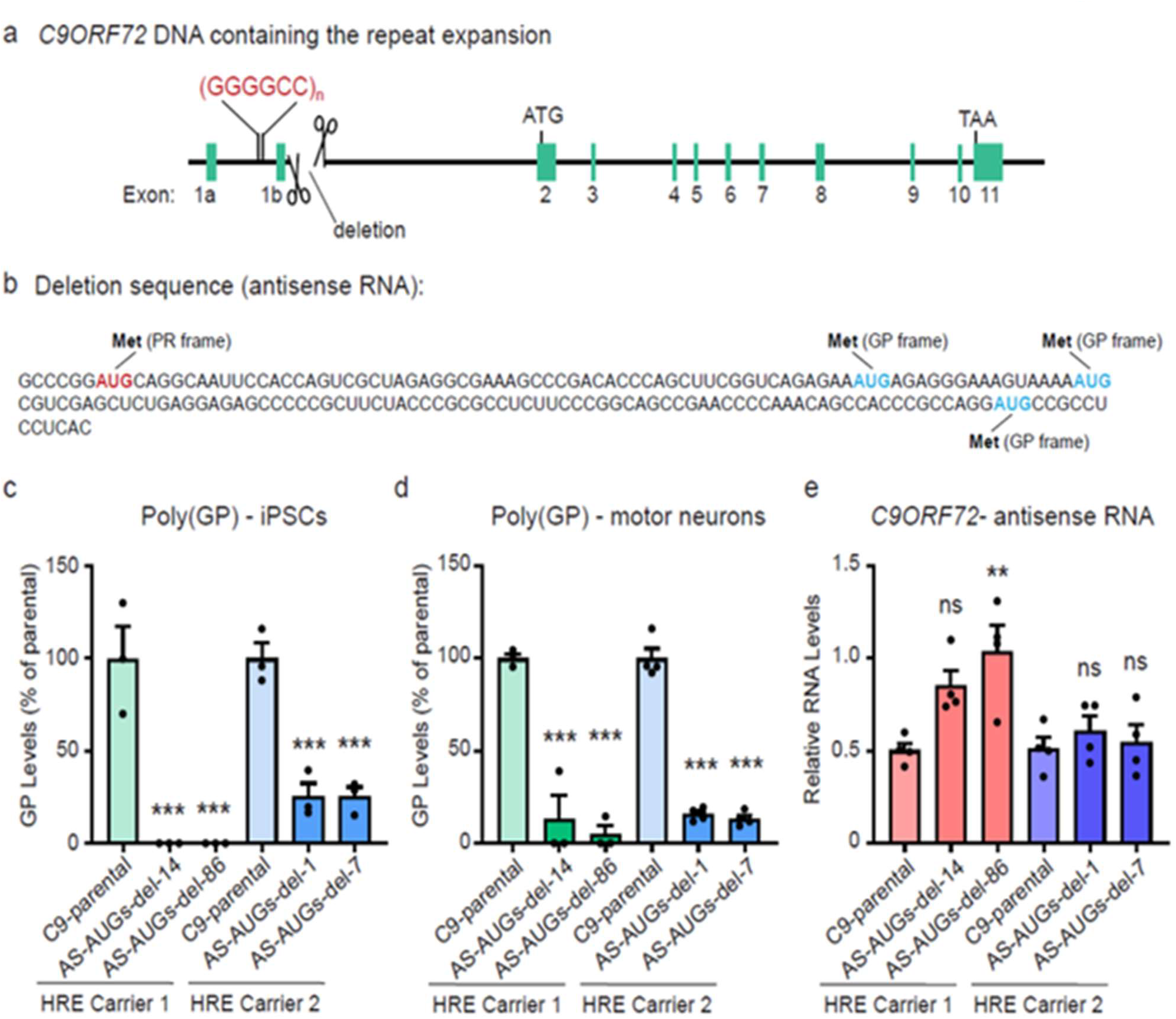
Deletion of the AUG codons region in the antisense repeat RNA reduces poly(GP) protein levels in *C9ORF72* iPSCs and motor neurons. **a** Schematic of the *C9ORF72* gene depicting the location of the 181-bp deletion generated by CRISPR-Cas9. **b** Antisense RNA nucleotide sequence of the deletion, highlighting the AUG codons in the poly(GP) and poly(PR) reading frames. **c, d** Poly(GP) protein levels in parental and antisense (AS)-AUGs-deletion iPSCs (**c**) and motor neurons (**d**) derived from two *C9ORF72* hexanucleotide repeat expansion (HRE) carriers. **e** Relative expression levels of *C9ORF72*-antisense RNA in parental and AS-AUGs-deletion motor neurons. Values are the mean ± s.e.m. of three to four independent differentiations. **p < 0.01, ***p < 0.001 (one-way ANOVA, Dunnett’s multiple comparisons test). ns: not significant. Met: methionine.

Two AS-AUGs-deletion lines from each parental *C9ORF72* iPSC line were selected for further analysis. All four iPSC lines contained the same 181-nucleotide homozygous deletion (Fig. 1b). We first determined the impact of this deletion on poly(GP) production by measuring its levels using a previously developed Meso Scale Discovery (MSD) assay^6^. Soluble poly(GP) was nearly undetectable in AS-AUGs-deletion iPSCs from HRE carrier 1 and more than 80% reduced in HRE carrier 2 (Fig 1c) compared to the respective parental line. Similar reductions in poly(GP) protein levels were observed after differentiation of all the iPSC lines into MNs (Fig. 1d). Importantly, AS RNA levels measured by quantitative RT-PCR did not decline (Fig. 1e). This result indicates that soluble poly(GP) levels decrease due to reduced translation rather than the absence or reduction of the RNA template in these lines. Examination of sense RNA translated soluble poly(GR) and poly(GA) showed that their levels were unaffected in AS-AUGs-deletion iPSCs or MNs (Fig. 2a–c). If the majority of poly(GP) synthesis occurred via sense repeat RNA translation then poly(GP) levels would be expected to remain unaffected in the AS-AUGs-deletion, contrary to our observations. Altogether, these results are consistent with the notion that the sequence containing the AUG start codons upstream of the C_4_G_2_ repeats in the AS repeat RNA is required to produce most of the soluble poly(GP) in *C9ORF72* neurons.

**Fig 2.**
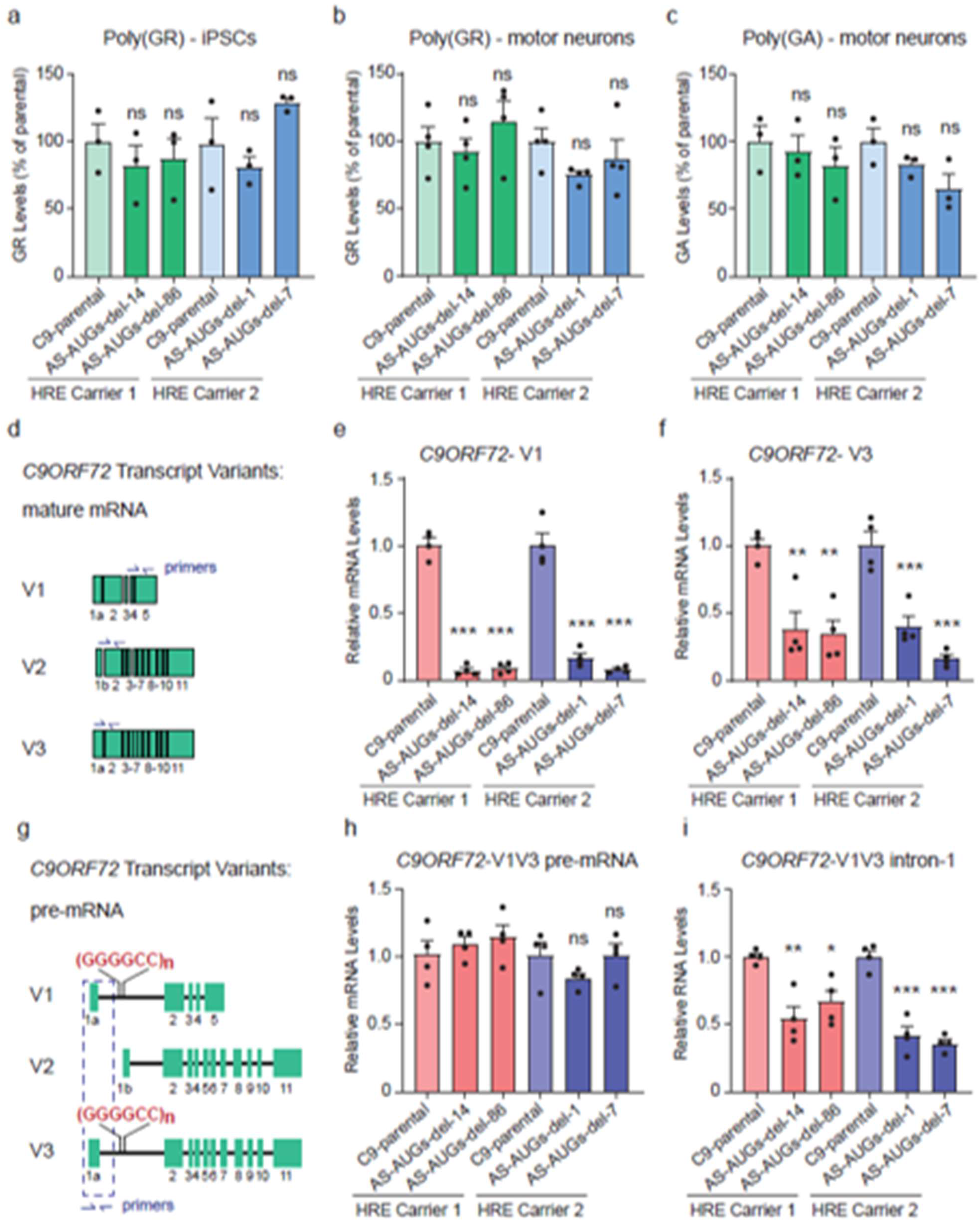
Poly(GR) and poly(GA) proteins are produced from the *C9ORF72* unspliced mRNA in AS-AUGs-deletion iPSC-derived motor neurons. **a–c** Poly(GR) protein levels in iPSCs (**a**) and motor neurons (**b**) and poly(GA) levels in motor neurons (**c**) measured with Meso Scale Discovery immunoassays. **d** Schematic representation of the spliced mRNA for V1, V2, and V3 of *C9ORF72*. The location of the primer sets used to detect each specific isoform is shown. **e, f** Relative expression levels of mature *C9ORF72* -V1 (**e**) and -V3 (**f**) in *C9ORF72* parental and AS-AUGs-deletion neurons. **g** Schematic representation of the pre-mRNA for the three *C9ORF72* transcript variants (V1, V2, and V3). The location of the primer set used to detect V1V3 pre-mRNA is shown. **h, i** Relative expression levels of *C9ORF72* pre-mRNA (**h**) and intron 1 (**i**) of both V1 and V3 in *C9ORF72* parental and AS-AUGs-deletion neurons. Values are the mean ± s.e.m. of three to four independent differentiations. *p < 0.5, **p < 0.01, ***p < 0.001 (one-way ANOVA, Dunnett’s multiple comparisons test). ns: not significant.

### Poly(GR) and poly(GA) proteins are produced from unspliced *C9ORF72* mRNAs

Since the AS-AUGs-deletion is close to the end of exon 1b (Fig. 1a), the first exon of *C9ORF72* variant 2 (V2), we examined whether splicing of V2 was affected. The forward primer used to detect mature V2 (Fig. 2d) spans the exon 1b-exon 2 junction and therefore only spliced V2 sequences can be amplified. In AS-AUGs-deletion motor neurons, mature V2 mRNA was undetectable by qRT-PCR, indicating that its splicing was blocked. Unexpectedly, the levels of mature mRNAs from both V1 and V3, which differ from V2 in that they incorporate an alternative exon to exon 1b (Fig. 2d, g), were also greatly reduced (Fig. 2e, f). How the deletion affects the splicing of V1 and V3 is not clear. Because mature mRNAs do not contain the repeat expansion, which is intronic and thus removed after splicing (Fig. 2d, g), we next examined whether pre-mRNA (unspliced mRNA) or intron-1 RNA levels were affected by the deletion. V1 and V3 pre-mRNA sequences are too similar to design isoform-specific primers, so we used a primer set that detects pre-mRNAs (mRNAs containing both exon 1a and intron-1, Fig. 2g) of both V1 and V3 (V1V3 pre-mRNA). In AS-AUGs-deletion motor neurons, pre-mRNA levels of V1 and V3 were equivalent to those of their respective parental neurons (Fig. 2h). Therefore, the AS-AUGs-deletion does not show any evidence of a reduction in the transcription of the repeat sense RNA. Under this experimental condition, the levels of intron 1 RNA, which accounts for both spliced and unspliced RNAs, was decreased by about 50% in AS-AUGs-deletion MNs, reflecting reduced splicing (Fig. 2i). These observations, together with the fact that the levels of soluble poly(GR) and poly(GA), which are translated from intronic sequences containing G_4_C_2_ repeats, are unaffected in AS-AUGs-deletion MNs, strongly supports the notion that V1/V3 pre-mRNA serves as the template for poly(GR) and poly(GA) DPR proteins in human neurons in this context (Fig. 2 a–c, h). Consistent with our findings, a recent study utilized *C9ORF72* HRE carrier derived MNs and estimated that at least 30% of transcripts containing exon 1a retain the repeat expansion^21^. The authors also showed that excision of only exon 1a by genome editing eliminated poly(GA) production^21^. Earlier studies reported the presence of RNAs containing unspliced intron-1 in cells and autopsy tissues of *C9ORF72* carriers and that unspliced RNA is a primary substrate for translation of RNAs containing the repeat expansion^3,19,25,29^.

### Antisense oligonucleotides targeting the sense *C9ORF72* repeat RNA reduce both sense and antisense RNAs

Our observation that most of the poly(GP) produced in *C9ORF72* neurons is dependent on the sequence containing the AUG start codons upstream of the C_4_G_2_ repeats in the AS RNA appears to conflict with recent reports. Administration of ASOs targeting the sense G_4_C_2_ repeat RNA (C9-ASOs) results in the loss of most poly(GP) in *C9ORF72* cells, a BAC transgenic mouse and *C9ORF72*-ALS/FTD repeat expansion carriers^5,9,12,27^. While these reports would seem to indicate that poly(GP) is mostly produced from the sense strand RNA, the levels of the AS repeat RNA were not determined in the studies. To investigate whether sense C9-ASOs affect the levels of AS RNA, we exposed MNs from HRE Carrier 1 to sense C9-ASOs for 6 days before assessing different *C9ORF72* RNAs (Fig. 3a). The two C9-ASOs used here, Ionis-672681 (first reported in Jiang et al 2016 and noted here as C9-ASO-1) and Afinersen (first reported in Tran et al 2022 and noted here as C9-ASO-2) target the same sequence in intron-1, which is only present in *C9ORF72*-V1 and -V3 RNAs (Fig. 3b). As expected, *C9ORF72* V1V3 intron-1 RNA levels were reduced by 50% in *C9ORF72* MNs treated with either of the C9-ASOs (Fig. 3c). The mature *C9ORF72*-V3 RNA, which does not contain the ASO target region, was less affected by the ASOs likely reflective of the quick splicing occurring in these cells (Fig. 3d). Mature *C9ORF72*-V2 (or its pre-mRNA) does not contain the ASO target region, and its levels were indeed unaffected by either ASO treatment (Fig 3e). Interestingly, *C9ORF72*-AS levels were decreased by more than 50% after ASO treatment in *C9ORF72* MNs (Fig. 3f). We obtained similar results using a second set of AS primers (first reported by Zu et al 2013) (Fig. 3g).

**Fig 3.**
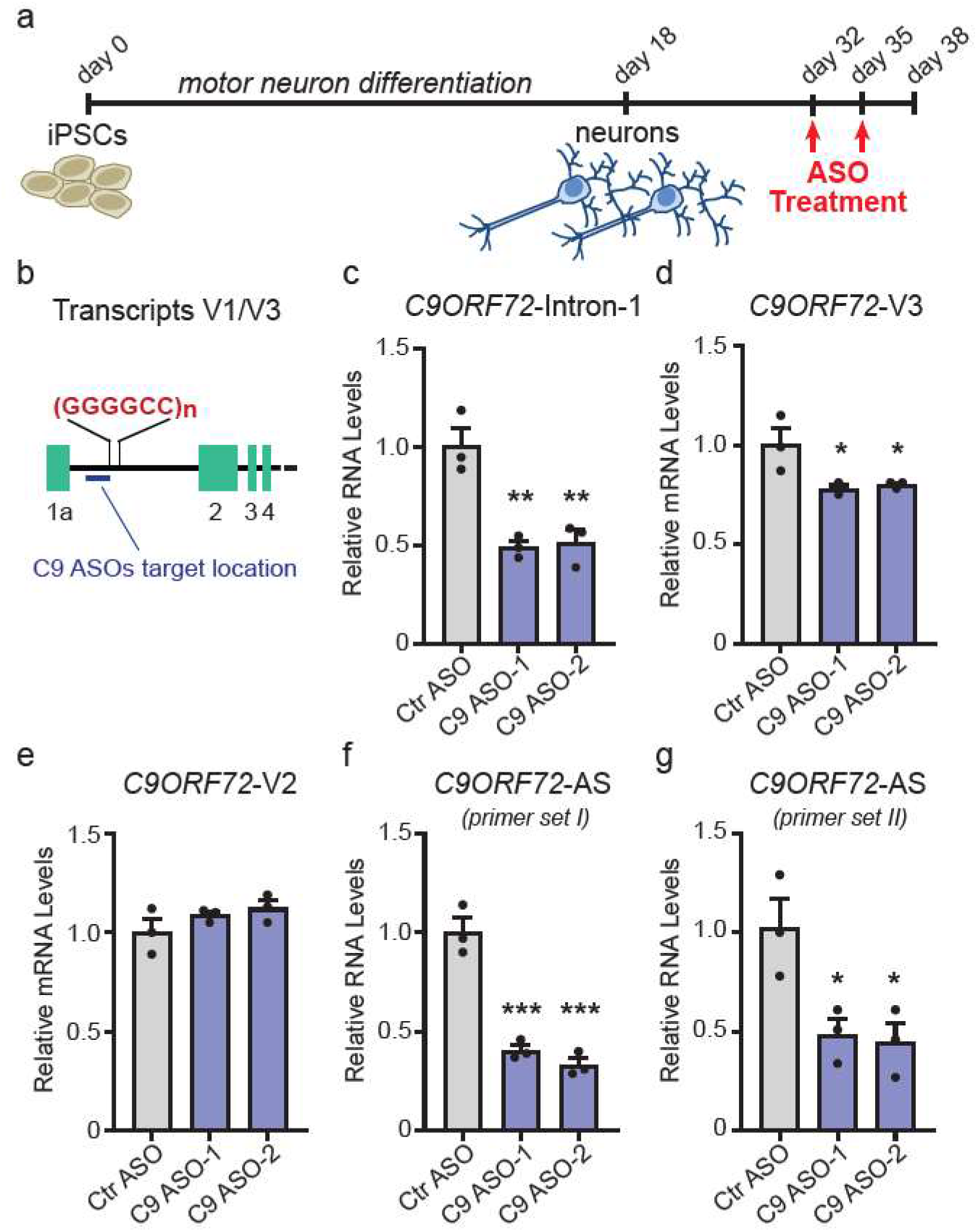
Antisense oligonucleotides (ASOs) targeting the *C9ORF72* sense repeat RNA reduce both sense and antisense RNAs. **a** Schematic representation of the timeline for the motor neuron differentiation and ASO treatment. **b** Diagram shows the region on V1 or V3 RNA targeted by the C9 ASOs. **c–g** Relative expression levels of *C9ORF72*-intron 1 (**c**) -V3 (**d**) -V2 (**e**) and antisense (AS) (**f, g**) in *C9ORF72* motor neurons treated twice with C9 ASO-1 or -2 for a total of six days. Two different sets of primers were used to measure AS RNA. Values are the mean ±s.e.m. of three independent differentiations. *p < 0.5, **p < 0.01, ***p < 0.001 (one-way ANOVA, Dunnett’s multiple comparisons test).

### C9-ASO targeting the sense repeat RNA decreases Poly(PR)

We then investigated whether the sense C9-ASO had a similar effect on the AS RNA of other cell types derived from *C9ORF72* HRE carriers. Previously, treatment of *C9ORF72* mutation carrier lymphoblast cell lines (LCLs) for ten days with a C9-ASO that targets the hexanucleotide G_4_C_2_ repeat sequence greatly reduced *C9ORF72*-V1, -V3 and poly(GP) levels^9^. To determine whether *C9ORF72* AS levels were also reduced under similar experimental conditions, we obtained LCLs from two previously characterized *C9ORF72* mutation carriers (ND10966, LCL1; ND11836, LCL2)^17^. As anticipated, C9-ASO-1 treatment greatly reduced *C9ORF72*-V3 and *C9ORF72* intron-1 RNA levels in both LCLs (Fig 4a, b). As observed in MNs, *C9ORF72*-AS RNA levels were also decreased to a similar extent (Fig. 4c). Altogether, the MN (Fig 3f, g) and LCL data support the surprising finding that sense C9-ASOs, in addition to decreasing the levels of sense RNAs, also reduce AS RNAs.

**Fig 4.**
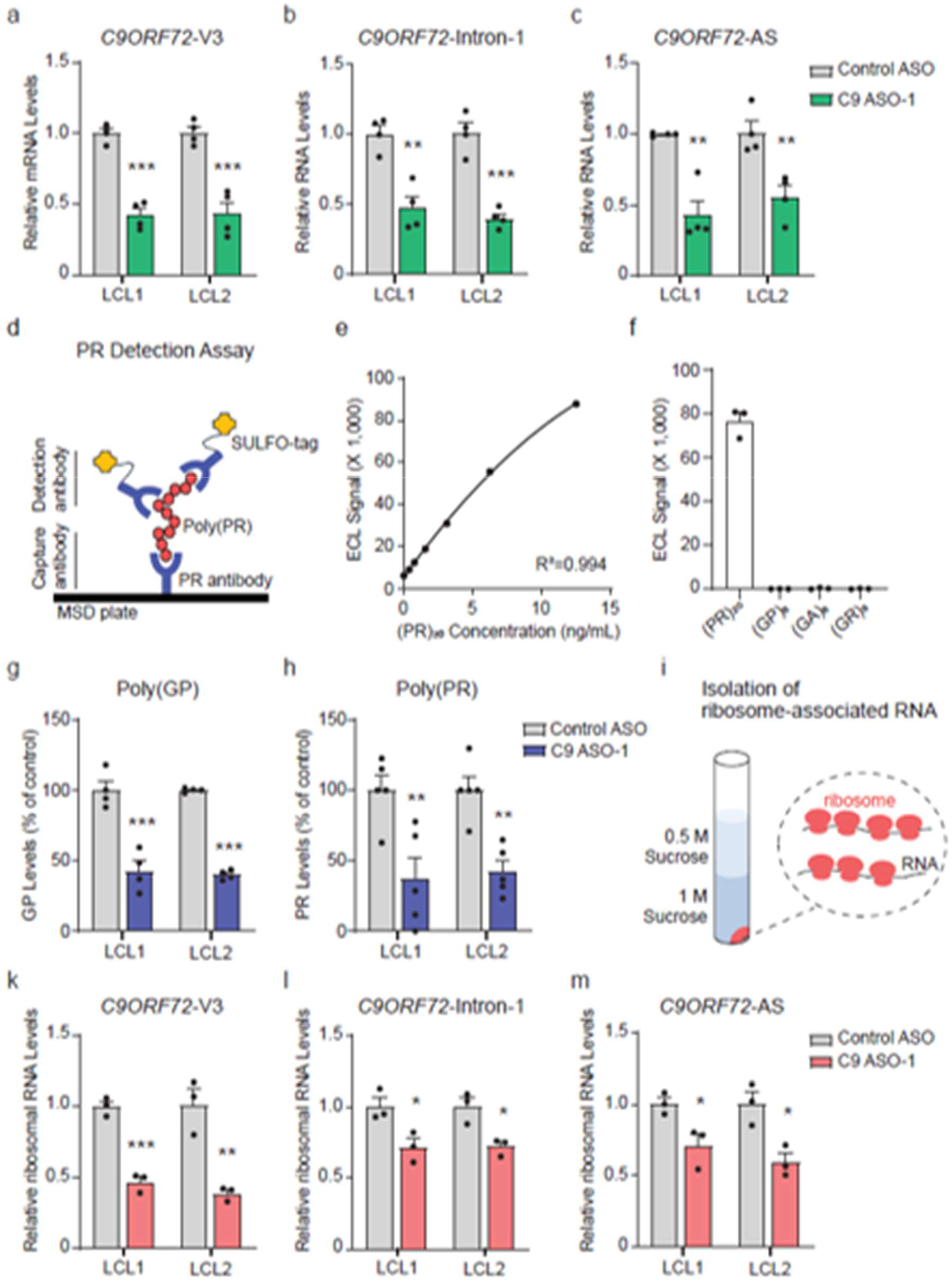
Poly(PR) protein synthesis decreases after treatment with ASO targeting the sense repeat RNA. **a–c** Relative expression levels of *C9ORF72*-V3 (**a**) -intron-1 (**b**) -antisense (AS) (**c**) in two *C9ORF72* mutation carrier lymphoblastoid cell lines (LCLs) treated with C9-ASO-1 for 10 days. **d** Diagram of the Meso Scale Discovery (MSD) assay developed to detect poly(PR) using a custom antibody. **e, f** Validation of the assay with a standard curve using increasing concentrations of a synthetic (PR)_20_ peptide (**e**) and a specificity test using (PR)_20_, (GP)_8_, (GA)_8_, (GR)_8_ peptides (**f**) showing that the assay only detects PR. **g, h** Poly(GP) (**g**) and poly(PR) (**h**) protein levels in two *C9ORF72* LCLs treated with C9-ASO-1 for 10 days. **i** Diagram of the sucrose gradient after ultracentrifugation to illustrate the pellet containing polysomes. **k–m** Relative expression levels of *C9ORF72*-V3 (**k**) -intron-1 (**l**) -AS (**m**) in the ribosome-associated RNA isolated from two *C9ORF72* LCLs treated with C9-ASO-1 for 10 days. Values are the mean ±s.e.m. of three to five independent cultures. *p < 0.5, **p < 0.01, ***p < 0.001 (two-tailed t test).

If the C9-ASOs decrease AS RNA levels, then the level of the DPR proteins that are translated from the AS RNA, such as poly(PR), should also be diminished. We therefore developed a specific MSD assay to detect poly(PR) that does not cross-react with poly(GP), poly(GA) or poly(GR) (Fig. 4d-f). As expected, the 10 day treatment with C9-ASO-1 reduced poly(GP) levels by more than half in the two *C9ORF72* LCLs (Fig. 4g). In addition, Poly(PR) levels were reduced by a similar extent (Fig. 4h), supporting the finding that the levels of the repeat AS RNA are reduced upon C9-ASO-1 treatment. We next investigated whether a decrease in the AS RNA translation could also contribute to the low levels of poly(PR) protein observed. Thus, we performed polysome analysis on LCLs treated with C9-ASO-1 to determine the translation levels of the different *C9ORF72* RNAs (Fig. 4i). Treatment with C9-ASO-1 led to a significant reduction in the translation of mature *C9ORF72*-V3 mRNA and sense RNAs that contained the intronic repeat expansion into protein (Fig. 4k, i). Similarly, there was a significant reduction in the amount of AS RNAs being translated into protein (Fig. 4m). Altogether, our data shows that sense targeting C9-ASOs not only decrease sense repeat RNAs, but also reduce AS repeat RNAs and their respective protein products. The results also indicate that the effect of the C9-ASOs on poly(GP) protein levels is not indicative of poly(GP) being generated from the sense RNAs, as initially speculated.

### Poly(PR) is produced by AUG-dependent translation of the antisense repeat RNA

The *C9ORF72* AS-AUGs-deletion iPSC lines were engineered not only to eliminate the three AUG start codons in the poly(GP) frame, but also the single AUG in the poly(PR) frame (Fig. 1a-b). Thus, we assessed whether this AUG plays a substantial role in the production of poly(PR) in human neurons, in a similar manner as the AUGs driving translation of the poly(GP) reading frame. Our customized MSD immunoassay for poly(PR) did not convincingly detect poly(PR) in human MNs derived from the two *C9ORF72* mutation carriers and were unable to determine whether the levels of poly(PR) were affected by the absence of the AUG start codon. As an alternative, we designed steric-blocking ASOs complementary to the region containing the AUG start codon in the poly(PR) frame and used these to treat the two *C9ORF72* LCLs following the same 10-day treatment protocol as was used for the C9-ASO-1 (Fig. 5a). The LCL-1 line was treated with three distinct ASOs targeting a sequence containing the AUG in the poly(PR) frame (ASOs 10-12), and one ASO targeting the AUG at the position -196 in the poly(GP) frame (ASO-13). This AUG at position -196 was previously shown to be the main initiation site that produces poly(GP) in a reporter system^4^. ASO-10 and -12 reduced poly(PR) protein levels by 40% and 30%, respectively. ASO-11 also decreased poly(PR) levels but did not reach statistical significance (Fig. 5b). The effect of ASO-10 was further confirmed in the LCL-2 line, in which a more than 60% reduction in poly(PR) protein was observed. Interestingly, ASO-13, which targets an AUG in the poly(GP) frame, also reduced the levels of poly(PR) protein in LCL-1, but not in LCL-2 (Fig. 5b). The effect of these steric-blocking ASOs on poly(GP) protein levels was also assessed. ASO-10 decreased poly(GP) by about 30% (LCL-1) and 50% (LCL-2), while ASO-12 displayed an approximately 65% reduction in LCL1 (Fig. 5c). However, ASO-13, which targets the AUG at position -196 in the poly(GP) reading frame, did not significantly reduce poly(GP) protein levels (Fig. 5c). One possible explanation for the observed lack of an effect of the ASO-13 is that there are two additional AUG start sites that are not targeted by the ASO in the poly(GP) frame that potentially drive the translation of poly(GP) in LCLs. The effect of the ASOs in these studies is likely acting at the level of translation, not transcription, of the repeat proteins because both the AS RNA (Fig. 5d) and the sense RNA (Fig. 5e, *C9ORF72*-Intron-1) levels are unaffected by treatment. Taken together, the data obtained with steric-blocking ASOs shows that the AUG codon in the poly(PR) frame drives repeat protein production through canonical translation of the AS repeat RNA.

**Fig 5.**
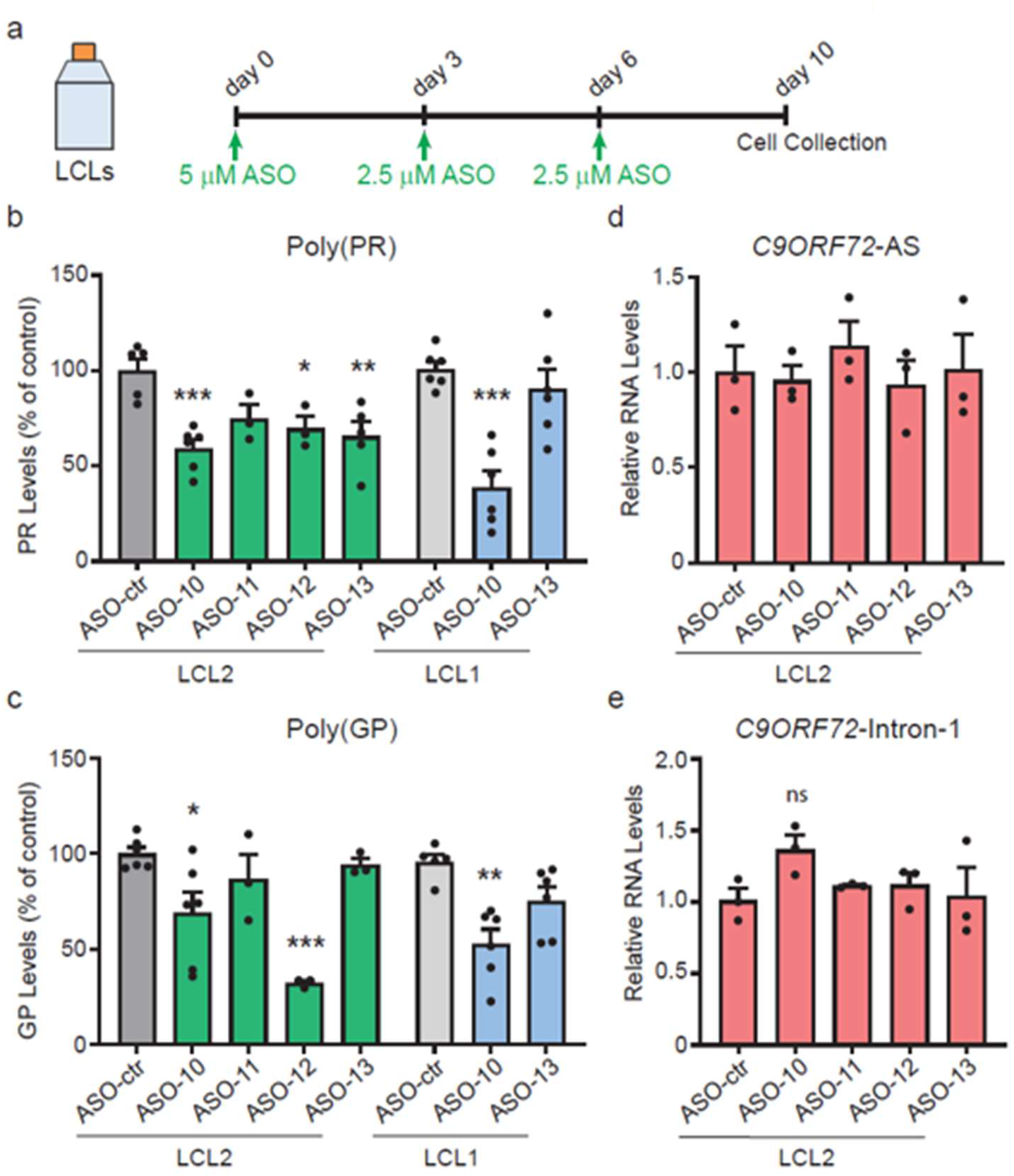
Antisense oligonucleotides (ASOs) steric-blocking the AUG start codon in the antisense repeat RNA suppress poly(PR) protein synthesis. **a** Schematic representation of the timeline for treatment of the *C9ORF72* mutation carrier lymphoblastoid cell lines (LCLs) with the ASOs steric-blocking the AUG start codons in the *C9ORF72* antisense repeat RNA. **b, c** Poly(PR) and poly(GP) protein levels in two *C9ORF72* LCLs treated with the steric-blocking ASOs. **d, e** Relative expression levels of *C9ORF72*-AS and -intron-1 in *C9ORF72* LCL1 treated with the steric-blocking ASOs for 10 days. Values are the mean ± s.e.m. of three to six independent cultures. *p < 0.5, **p < 0.01, ***p < 0.001 (one-way ANOVA, Dunnett’s multiple comparisons test). ns: not significant.

## Discussion

The mechanism by which DPR proteins are synthesized from RNAs harboring repeat expansions in *C9ORF72* mutation carrier cells is not fully elucidated. Multiple translation mechanisms likely play a role in the production of these proteins. Our results highlight the important influence of the sequences preceding the hexanucleotide repeat expansion for the synthesis of poly(GP) and poly(PR) in *C9ORF72* mutation carriers MNs and LCLs. We found that poly(GP) and poly(PR) are produced from the AS repeat RNA through a translation mechanism that requires an AUG start codon. In the poly(GP) reading frame, there are 3 putative AUG initiation sites (at 212, 194 and 113 nucleotides upstream of the repeats), which are not interrupted by in-frame termination codons in between the AUGs and the C_4_G_2_ repeats. Any or all of these AUGs could initiate poly(GP) protein production. Boivin et al (2020) analyzed by mass spectrometry the poly(GP) protein generated via expression of the anti-sense sequence in cells and found that the AUG start codon located 194 nucleotides upstream of the C_4_G_2_ repeats was used to initiate poly(GP) protein production, while deletion of the AS sequence containing this AUG codon precluded poly(GP) protein production^4^. Intriguingly, deletion of the sequence containing the 3 AUGs in MNs decreased the production of the majority of poly(GP), whereas the steric blocking ASO targeting only the AUG codon located at position 194 (ASO-13) did not have a significant effect on poly(GP) protein levels. ASO-13 was able to decrease poly(PR) levels, an indication that the ASO does engage its intended target sequence. Therefore, we surmised that the prevention of poly(GP) protein production requires disruption of more than one AUG initiation codon. In line with this finding, Sonobe et al used a luciferase reporter to show that while the AUG at 194 appears to be the main start codon for poly(GP) protein production, two other AUG codons (at position 212 and 113 from the repeats) can function as alternative translation initiation sites when the AUG at 194 was mutated^24^. Thus, a therapeutic strategy aimed at reducing poly(GP) protein would likely need to address prevention of translation initiation from all 3 sites. For reducing poly(PR), the AUG initiation site located at position 273 from the repeat expansion would serve as a potential site of intervention. Interestingly, targeting poly(PR) production also partially reduced poly(GP) protein, raising the possibility that a single poly(PR) ASO treatment may more broadly impact the synthesis of additional DPR proteins.

An important implication revealed in this work is that DPR proteins appear to have unique N-termini in addition to distinct C-termini^31^. This has relevance since the particular amino acid composition of these proteins can affect their structure, localization, interaction partners and ultimately toxic potential. In line with this, the inclusion of the native C-terminal sequence was found to reduce toxicity and change the subcellular localization of DPR proteins^11^. Future studies evaluating the toxic potential of DPR proteins need to assess their native sequences to ensure findings are relevant for the endogenous disease context.

Another major insight of this work helps to further reconcile whether sense or AS repeat RNAs are the most prevalent source of poly(GP). Our deletion data clearly shows that in *C9ORF72* derived MNs the major source of poly(GP) protein is the AS repeat RNA. This is supported by a recent study that showed that CRISPR/Cas9 mediated excision of exon 1a in *C9ORF72* MNs eliminated sense-dependent poly(GA) but did not significantly impact poly(GP) protein^21^. In this study, most of the AS RNA remained while abolishing sense RNA, suggesting that poly(GP) protein is produced from the AS direction^21^. Our observations and those of Sachdev et al are also consistent with *C9ORF72* expansion carrier autopsy tissue studies which showed that about 80% of poly(GP) protein is generated from AS repeat RNAs^31^. These findings are contrasted by reports showing ASOs that target sense *C9ORF72* RNA greatly reduce poly(GP) protein in cells and cerebrospinal fluid from expansion carriers^9,27^. However, it is important to note that these studies did not assess AS RNA levels. We found that two C9-ASOs that were previously administered to *C9ORF72* HRE carriers after disease onset^5,27^ decreased both sense and AS RNAs in HRE MNs. Consistent with these findings, the protein levels of the AS-dependent DPRs, poly(GP) and poly(PR), were also found to be greatly reduced. Our current hypothesis is that the effect of the sense C9-ASOs on the AS RNA is related to a local inhibition of transcriptional activity triggered by the ASO. Previous studies have shown that intron-targeted ASOs like the C9-ASOs used here cause transcription termination by reducing the density of RNA Polymerase II downstream of the ASO cleavage site^14,16^. In that study, the greatest impact on transcription was observed more than 20 kb downstream of the site of the ASO-directed RNA cleavage^14^. Another possibility is that C9-ASOs may alter local transcriptional activity by modulating the methylation state of chromatin histones, as was recently reported for the ASO used as a therapy for Spinal Muscular Atrophy, nusinersen^18^. While these unintended on-target effects may present challenges for the use of some ASOs as therapeutics, they may be advantageous in the case of *C9ORF72*. One ASO acting to reduce or eliminate both sense and AS RNA-driven toxicities may be more efficacious than providing a therapeutic mechanism targeting a single source of toxic species. A recent ASO clinical trial targeting sense repeat RNA unfortunately did not provide a desired outcome for people with *C9ORF72* ALS/FTD though it remains unclear whether this clinical outcome resulted from insufficient target engagement due to the timing of therapeutic administration and/or highlights a role for AS repeat pathologies, such as poly(PR) toxicity in C9ORF72 disease^5,28^. Given that we observed a poly(PR) reduction using sense targeting ASOs, it would be interesting to test patient samples from the above mentioned clinical trials for changes to poly(PR). Nevertheless, the current study provides evidence that employing a sense ASO strategy is capable of reducing poly(PR) and poly(GP) protein levels in *C9ORF72* HRE carrier derived cell types and can potentially serve as the basis for adapting strategies aimed at reducing AS repeat RNA products as potential therapeutics for *C9ORF72*-dependent ALS/FTD.

## Methods

### Generation of *C9ORF72* antisense-AUGs-deletion iPSC lines

The CRISPR-Cas9 system was used to create a deletion in the first intron of *C9ORF72*, 3’ to the G_4_C_2_ repeats at ALSTEM (Richmond, CA). In brief, the Amaxa nucleofector II (program B-016) was used to transfect iPSCs with two guide RNAs: CGGTGGCGAGTGGGTGAGTG and ATTGCCTGCATCCGGGCCCC. Nucleofected cells were treated with puromycin, dissociated into single cells, placed in 96-well plates, cultured for 14 days, and expanded. Genomic DNA from each clone was extracted with a genomic extraction kit (Zymo Research). Clones with the desired homozygous deletion were identified with a PCR amplification assay, and the PCR products were sent for sequencing. At least two iPSC lines containing the expected deletion from each *C9ORF72* expansion carrier were expanded and collected to isolate genomic DNA. The region of interest was amplified by PCR, and the products were sent for sequencing to confirm the identity of each clone.

### Motor neuron cultures

Motor neurons were differentiated from iPSCs as described^1^. Briefly, iPSC colonies were seeded on Matrigel-coated wells in mTeSR1 medium (StemCell Technologies); 24 h later, the medium was changed to neuroepithelial progenitor (NEP) medium consisting of 1:1 DMEM/F12:Neurobasal, 0.5x N2, 0.5x B27, 0.1 mM ascorbic acid (Sigma-Aldrich), 1x Glutamax, 3 μM CHIR99021 (StemCell Technologies), 2 μM DMH1 (StemCell Technologies), and 2 μM SB431542 (Stemgent) and replaced every other day for 6 days. Progenitor colonies were dissociated with Accutase, seeded 1:6 on Matrigel-coated wells, and cultured in NEP medium containing 0.1 μM retinoic acid and 0.5 μM purmorphamine for 6 days; the medium was replaced every other day. Motor neuron progenitors were lifted, cultured in suspension for 6 days in the absence of CHIR99021, DMH1, and SB431542, and dissociated to single cells with Accutase. Cells were seeded on poly-lysine/laminin-coated wells in motor neuron medium (1:1 DMEM/F12:Neurobasal, 0.5x N2, 0.5x B27, 0.1 mM ascorbic acid, 1x Glutamax, 10 ng/mL BDNF, 10 ng/ml GDNF, 1 μg/ml laminin, 0.1 μM compound E, 0.5 μM retinoic acid and 0.1 μM purmorphamine) for up to 4 weeks. This protocol generated a culture with >90% ChAT^+^ neurons. Cultures were prepared from two sets of lines derived from two C9ORF72 expansion carriers and respective no-repeats isogenic controls. The no-repeats iPSC lines were first reported in Lopez-Gonzalez et al., 2019. All experiments were done with neurons derived from 3–4 independent differentiations.

### Lymphoblast cell lines (LCLs) cultures

Lymphoblast cell lines ND10966, ND11836 and ND16183 were obtained from the Coriell NINDS Repository (Camden, NJ, USA). Lines ND10966 and ND11836 were derived from a *C9ORF72* repeat expansion carrier and denoted LCL1 and LCL2. Line ND16183, derived from a non-expanded ALS patient, was used to set the background for the DPR measurements. All lines were maintained in RPMI-1640 containing 15 % fetal bovine serum at 37 °C with 5% CO2.

### Antisense oligonucleotide (ASO) Treatment

Two-week-old motor neurons were treated twice with 5μM of previously described ASOs that target the region 5 ‘to the *C9ORF72* hexanucleotide G4C2 repeat sequence for a total of six days (Fig. 3a). The C9 ASO developed by Ionis Pharmaceuticals (ISIS# 672681, Jiang et al., 2016) was denoted C9-ASO-1 and the C9-ASO developed by J Watts and R Brown laboratories (Afinersen, Tran et al., 2022) was denoted C9-ASO-2. As a control ASO, we used an ASO (ISIS# 676630, Jiang el al., 2016) also generated by Ionis Pharmaceuticals.

Lymphoblastoid cell lines (LCLs) were treated with C9-ASO-1 following a protocol described by Gendron et al^9^. Briefly, LCLs were seeded at 4.5 × 10^6^ cells per T25 flask and treated with 5 μM control ASO or C9-ASO-1. Three and six days later, cells were seeded and treated again with 2.5 μM control or C9-ASOs. Cells were harvested ten days after seeding.

For the experiments with the steric-blocking ASOS, LCLs were treated with the same protocol as described for C9-ASO-1. ASOs 10-13 were designed by S Almeida and purchased from Integrated DNA Technologies (IDT). Control ASO was also obtained from IDT. All ASOs are full 2’-O-methoxyethyl (MOE), endonuclease resistant and designed to sterically block the region around the AUG codon in the PR frame (ASOs 10-12) or the AUG codon at position -196 in the GP frame on the AS RNA (ASO-13).

### Measurement of poly(GR) and poly(GP)

Poly(GR) and poly(GP) were measured with an MSD immunoassay as described^6^. Briefly, neurons were lysed in ice-cold RIPA buffer (Thermo Fisher Scientific) containing a cocktail of protease and phosphatase inhibitors (Thermo Fisher Scientific), sonicated on ice at a 20% pulse rate for 15 s and centrifuged at 16,000 *g* for 20 min at 4°C. The protein content of supernatants was determined with the Bio-Rad Protein assay reagent (Bio-Rad). Neurons or iPSC samples were loaded on a 96-well single-spot plate (MSD; Cat. No. L45XA) pre-coated with a custom-made polyclonal rabbit anti-(GR)_8_ or anti-(GP)_8_ antibodies (1 µg/ml, Covance) and tested in duplicate wells. Serial dilutions of recombinant (GR)_8_ or (GP)_8_ peptide in 1% BSA-TBST (Tris-buffered saline (TBS)-Tween20 (0.05%)) were used to prepare the standard curve. The detection antibodies were anti-(GR)_8_ or anti-(GP)_8_ antibodies previously tagged with GOLD SULFO (GOLD SULFO-TAG NHS-Ester Conjugation Pack, Cat. No. R31AA, MSD) at a concentration of 0.5 µg/ml. Response signals from the assay plate were acquired with a QuickPlex SQ120 instrument (MSD). For background correction, electrochemiluminescence (ECL) signal values from neuron/iPSC/LCLs samples lacking repeats were subtracted from the corresponding test samples.

### Measurement of poly(PR)

Poly(PR) assay was performed like the above section, using a custom-made, affinity-purified rabbit polyclonal PR antibody raised against a (PR)20 peptide (Covance) and following a protocol provided by MSD. Briefly, 96-well single-spot streptavidin plates (MSD; Cat. No. L45SA-1) were blocked with Blocker A, coated with biotinylated anti-PR antibody at a concentration of 0.5 μg/ml and incubated overnight at 4 °C. The next day, the plates were washed three times with TBST, loaded with samples in duplicate wells and incubated for 2 h at room temperature on a shaking platform. The dilutions for the standard curve were prepared with recombinant (PR)_20_ as described above. After three washes, SULFO-tag conjugated detection PR antibody was added to the plates (0.5 μg/ml), incubated for 2 h at room temperature on a shaking platform, and washed again. Lastly, the MSD-Read buffer (2×) was added, and the plates were read on the MSD instrument. Background correction was performed against LCLs samples lacking repeats and the ECL values were presented as relative to LCLs treated with control ASO.

### Measurement of poly(GA)

Neuron pellets were thawed on ice in approximately 120 µl of lysis buffer (1x TBS, pH 7.4, 1 mM EDTA, 1% Triton X-100) with protease inhibitor cocktail (cOmplete, Sigma-Aldrich), vortexed, and incubated at 4°C for 15 min to fully lyse the pellet. Lysed cells were centrifuged at 14,000 g for 20 min at 4°C. Total protein concentration of the remaining supernatant was determined with the BCA protein assay (Thermo Scientific). Poly(GA) content was measured with an MSD sandwich immunoassay. In this assay, the human/murine chimeric form of anti-GA antibody chGA3 is used for capture, and human anti-GA antibody GA4 and a SULFO-tagged anti-human secondary antibody are used for detection. Poly(GA) concentrations were interpolated from the standard curve using 60X-GA expressed in HEK 293 cells and expressed as ng/mg total protein. For background correction, values from neuron samples lacking repeats were subtracted from the corresponding test samples.

### Polysome isolation

Polysomes were isolated from LCLs treated with C9 ASO-1 as described^29^. Briefly, LCLs were treated with 100 μg/mL cycloheximide (CHX) for 1 min, washed with PBS containing CHX and lysed in ice-cold polysome buffer (20 mM Tris–HCl pH 7.4, 5 mM MgCl2, 100 mM KCl, 1mM DTT, 100 μg/mL CHX, 0.3% NP-40, 40 U/mL RNase inhibitor and 1 tablet EDTA-free protease inhibitor) for 15 min. The lysates were centrifuged at 2,000g for 10 min and then at 20,000g for 10 min at 4°C. The final supernatants were further centrifuged at 80,000 rpm for 3 h (OptimaMax-TL, TLA110 rotor) through 0.5 M and 1 M sucrose in polysome buffer. The RNA was extracted from the polysome pellet with Qiazol lysis reagent and the miRNeasy kit, treated with DNase I (Qiagen) and quantified by RT-qPCR.

### Gene expression

Total RNA was extracted with the Qiagen RNeasy kit and treated with DNase I. RNA (1–2 μg) was reverse transcribed into cDNA with random hexamers or *C9ORF72* antisense–specific reverse primer using the High Capacity cDNA kit (Thermo Fisher Scientific). Quantitative PCR was done with an Applied Biosystems Quant Studio 3 system and SYBR Select Master Mix (Thermo Fisher Scientific). Ct values for each sample and gene were normalized to cyclophilin or actin beta. The relative expression of each target gene was determined with the 2^−ΔΔCt^ method. The following primers were used for qPCR: *C9ORF72* AS–specific RT rev (Set I): 5’ cgactggagcacgaggacactgagggacaagggatggggatc 3’; AS fwd: 5’ ctctcagtacccgaggctc 3’; rev: 5’cgactggagcacgaggacactga 3’;

V1 fwd: 5’gagaatggaagatcagggtca 3’; rev: 5’gtatctgcttcatccagcttt 3’;

V2 fwd: 5 ‘cggtggcgagtggatatct 3’; rev: 5 ‘gcccaaatgtgccttactct 3’;

V3 fwd: 5’gggtctagcaagagcaggtg 3’; rev: 5’agcccaaatgtgccttactc 3’;

V1V3 pre-mRNA fwd: 5’ tcaaacagcgacaagttccg 3’; rev: 5’ ggagagagggtgggaaaaac 3’;

intron-1 fwd: 5’ccccactacttgctctcaca 3’; rev: 5’ctacaggctgcggttgtttc 3’.

### Statistical Analysis

Plots and statistical analyses were performed with GraphPad Prism 10 software. The type of error bar and statistical test used in each graph is specified in the figure legends. Correction for multiple comparison was done using Dunnett’s statistical test. Differences were considered significant if P < 0.05. The number of biological replicates is presented in each figure legend.

## Acknowledgements

We are grateful to Rigel Chan (UMass Chan) for helping with the request of LCL lines, Heleen van ‘t Spijker (UMass Chan) for providing the polysome isolation protocol and Gopinath Krishnan (UMass Chan) for discussions related to the biotin MSD plates setup. S.A. is supported by the NIH (R21NS119952, R21NS112766 and 1R21AG085076). We would like to also thank Fen-Biao Gao (UMass Chan) for kindly sharing his laboratory space and equipment and providing the polyGP antibody. F.B.G. was supported by R37NS057553 and R01NS101986. The poly(GA) antibodies used in the MSD assay were developed by Neuroimmune AG (Zurich, Switzerland).

## Author contributions

Y.G. and M.W.K. performed the MSD assay for polyGA. J.W. and P.J-N. provided the C9-ASOs. S.A. planned the study and performed and analyzed all remaining experiments. S.A. wrote the manuscript with input from all authors.

